# On the impact of the pangenome and annotation discrepancies while building protein sequence databases for bacteria proteogenomics

**DOI:** 10.1101/378117

**Authors:** K.C.T. Machado, S. Fortuin, G.G. Tomazella, A.F. Fonseca, R. Warren, H.G. Wiker, S.J. de Souza, G.A. de Souza

## Abstract

In proteomics, peptide information within mass spectrometry data from a specific organism sample is routinely challenged against a protein sequence database that best represent such organism. However, if the species/strain in the sample is unknown or poorly genetically characterized, it becomes challenging to determine a database which can represent such sample. Building customized protein sequence databases merging multiple strains for a given species has become a strategy to overcome such restrictions. However, as more genetic information is publicly available and interesting genetic features such as the existence of pan- and core genes within a species are revealed, we questioned how efficient such merging strategies are to report relevant information. To test this assumption, we constructed databases containing conserved and unique sequences for ten different species. Features that are relevant for probabilistic-based protein identification by proteomics were then monitored. As expected, increase in database complexity correlates with pangenomic complexity. However, *Mycobacterium tuberculosis* and *Bortedella pertusis* generated very complex databases even having low pangenomic complexity or no pangenome at all. This suggests that discrepancies in gene annotation is higher than average between strains of those species. We further tested database performance by using mass spectrometry data from eight clinical strains from *Mycobacterium tuberculosis*, and from two published datasets from *Staphylococcus aureus.* We show that by using an approach where database size is controlled by removing repeated identical tryptic sequences across strains/species, computational time can be reduced drastically as database complexity increases.

## Introduction

While genome annotation approaches undertook considerable advances in the past years(Bocs et al., 2003;Davidsen et al., 2010;Vallenet et al., 2013), correct gene prediction is still challenging even for prokaryotes. Particularly, the correct assignment of open reading frames (ORFs), the presence and classification of pseudogenes anddifferences in translational starting site (TSS)(de Souza et al., 2008;de Souza et al., 2009;Cuklina et al., 2016) are the main sources of variability. Consequently, for any given strain or species, its predicted protein sequence information will normally contain inaccuracies that will interfere with peptide identification in mass spectrometry (MS)-based proteomics. The use of accurate protein sequence databases is a key step in most proteomic approaches, and this is particularly critical for studies involving samples containing strains with little to none genetic information or samples containing multiple strains (metaproteomics) (Tanca et al., 2013).

In such cases when the establishment of a gold-standard annotation that can represent the sample under investigation is difficult, a viable alternative is to construct customized protein sequence databases which are then inspected against peptide sequence data collected by MS (Nesvizhskii, 2014). This has been largely employed in proteogenomics, where “novel” sequences are inserted in the customized database and, if identified, are further used to validate and confirm proposed gene models and other genetic polymorphisms (for recent reviews see (Renuse et al., 2011;Ruggles et al., 2017)). Database customization is often achieved using two different strategies: i) if the genomic information of the prokaryote under study is known, a 6-frame translation is possible (Fermin et al., 2006;Baerenfaller et al., 2008); ii) if genomic information is unknown, an alternative is to construct a database by merging *ab initio* gene predictions from related strains of the same species, taking into consideration variations caused by SNPs, indels, divergent TSS choice, among others (de Souza et al., 2010;Omasits et al., 2017). These approaches are not mutually exclusive, as gene annotation from related strains can be used to further optimize 6-frame translation approaches (Castellana et al., 2008).

However, peptide identification in MS-based proteomics is often performed through probabilistic calculations between the observed peptide fragmentation pattern and theoretical data from a sequence database. Therefore, database size will: i) alter the search space and consequently the probabilistic calculations performed during peptide identification (Nesvizhskii, 2010); ii) make protein inference more difficult, especially in multistrain databases (Nesvizhskii and Aebersold, 2005); and iii) demand more computational power due to the handling of larger files. Therefore, building databases using either 6-frame translations or sequence merging approaches which are larger than regular can become, at some point, detrimental for the proteomic analysis (Blakeley et al., 2012).

Such issue might as well contribute differently depending on the species under study, how much genomic information is available (number of strains with genome sequenced) and genomic features particular to it. For example, with few sequenced strains at the time, we previously observed that the size of such merged database for *Mycobacterium tuberculosis* (Mtb) complex was not heavily incremented as more strains were considered (de Souza et al., 2011). Each new strain added to the database contributed with only few genetic polymorphisms when compared to the reference H37Rv strain. However, the current 65 Mtb strains with complete genome sequenced demonstrated the existence of a pangenome in this species, i.e., the genome of all strains of a given species contains only a set of genes from a larger pool of accessory genes for that species (Rasko et al., 2008;McInerney et al., 2017). It is then critical to address the impact of such approach in the size of customized databases used in MS-based proteomics, as well as in the performance for peptide identification.

Many reports have demonstrated that database size could be controlled by creating protein entries where identical peptides between homologue proteins are reported once and only unique polymorphic tryptic peptides are inserted when merging sequences (Schandorff et al., 2007;Omasits et al., 2017;Liao et al., 2018). It is unclear if genomic features such the presence of a pangenome could impact database customization of specific species. We then decided to test this by creating protein sequence databases for ten species using strains with complete genome sequences and a modified version of MSMSpdbb (de Souza et al., 2010). Database parameters such as the number of entries and unique peptide sequences for each species using increasing number of strains per round were then quantified. As expected, species with large pangenomes such as *Escherichia coli* showed the larger increase in database size per number of strains used, and database size in general correlated with pangenome size. Finally we performed proteomic identification and computational performance oftwo different datasets from: i) eight clinical strains of Mtb using a database with 65 complete strain genomes; and ii) two *S. aureus* datasets (Depke et al., 2015;Hoegl et al., 2018) using a database with 194 complete strain genomes.

## Experimental Procedures

### Species selection

Ten bacterial species were selected for database construction: *Acinetobacter baumannii* (89), *Bordetella pertusis* (348), *Campylobacter jejuni* (114), *Chlamydia trachomatis* (82), *Escherichia coli* (425), *Listeria monocytogenes* (148), *Mycobacterium tuberculosis* (65), *Pseudomonas aeruginosa* (105) and *Staphylococcus aereus* (194). *Burkholderia mallei* and *Burkholderia pseudomallei* (109) are considered genetically very similar (Godoy et al., 2003;Song et al., 2010) and were analyzed together. Number in parenthesis represents strains with complete genome sequenced according to Genbank (Benson et al., 2013) in August 2017. Protein sequences were obtained using the assembly file available at ftp://ftp.ncbi.nih.gov/genomes/genbank/bacteria/.

### Script design and availability

We redesigned MSMSpdbb (de Souza et al., 2010) in PERL (version 5.24.1) and is present as two modules: all_fasta.pl provides the sequence alignment and creates the outputs with unique entries and homologues; create_bd.pl process the homologues output, and create the final database and the log file. In house BLAST installation (version 2.7.1 or later) is required. All script outputs are saved as txt files in the data folder. Rand.pl is also available to reproduce the experimental design described below.

### Data processing

Even though MSMSpdbb is already published, since we redesigned how homology is defined (using a Bidirectional Best Hit rather than a Cluster BLAST approach), we provide below further information about the method. To assign gene homology, our script initially gathers the protein sequence data from all strains and then performs pair wise alignments using BLASTP (Altschul et al., 1990) through a Bidirectional Best Hit (BHH) method (Overbeek et al., 1999). The script performs in a manner that the strain with most number of protein entries is initially selected as the query dataset and consequently aligned to the remaining strains sequence datasets (subjects). Two sequences from different strains will be defined as homologues if: i) they are the best hit possible for all alignments performed; ii) sequence identity is equal or higher than 50%; iii) sequence similarity is equal or higher than 70%; and iv) sequence coverage is equal or higher than 70%. These have to be correlated on both directions (BHH) of the alignment.

When homologues are defined between query and subject strains, our script will proceed by indexing homologue IDs for later characterizations of polymorphisms. It will then define “unique” protein sequences from the query dataset (i.e., a sequence without a defined homologue in any of the subject strains), and add it to the partial version of the final database output (see Figure 1). In parallel, all entries from the subject strains which were properly aligned to a query sequence will be removed from the respective strains datasets. A new round of alignment will be performed after excluding the query strain and selecting one of the previous subject datasets as a new query strain. This will be performed successively until all strains are used as query. At this point, the script will had then defined all “unique” protein entries from all datasets which did not aligned to any other annotated sequence under the selected parameters.

**Figure 1.**
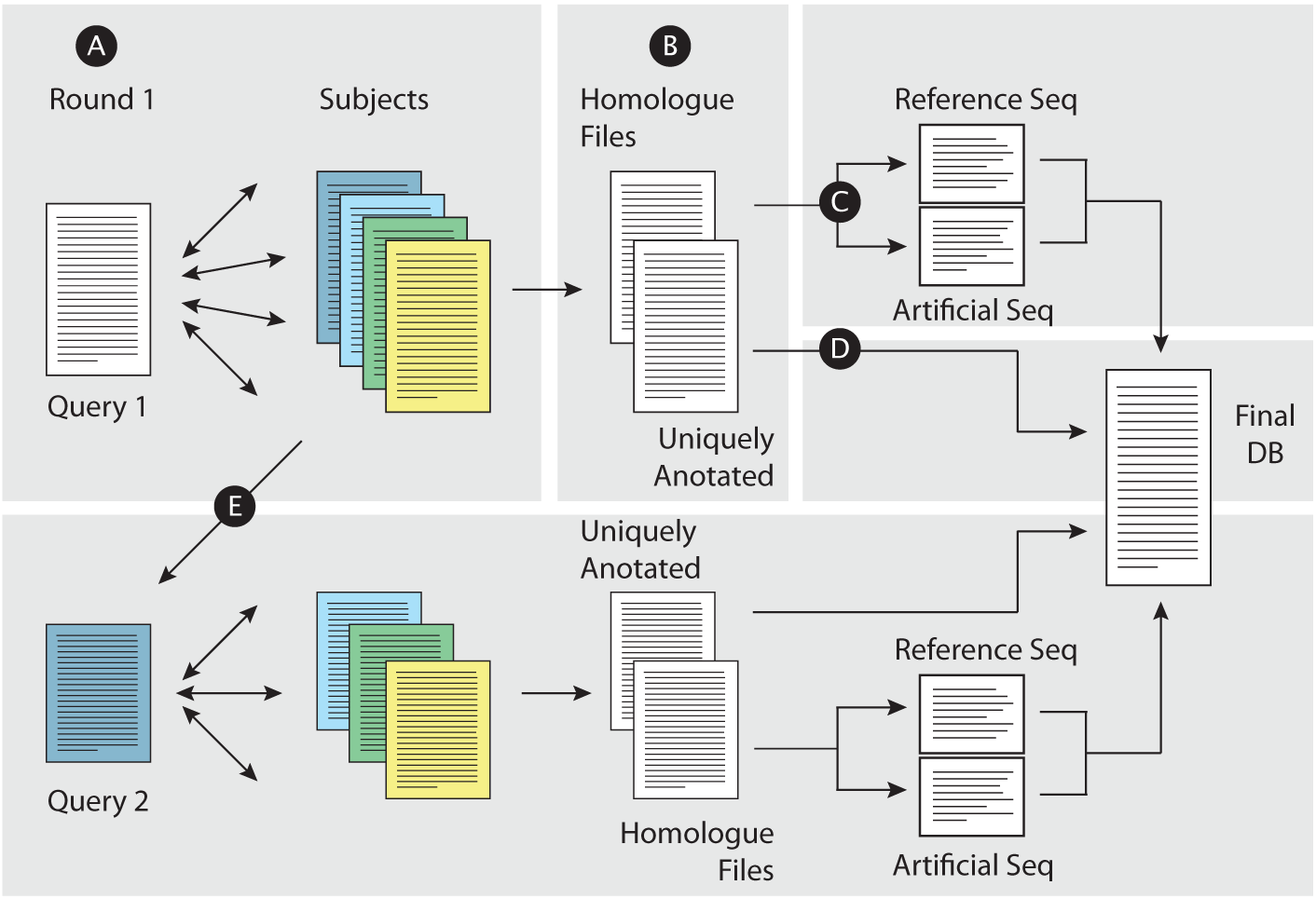
Building concatenated databases from related strains. (A) For a given set of fasta files, a Query file is selected and BLASTP is performed against the remaining files. A bidirectional approach is performed. (B) Uniquely annotated entries in the Query are defined and moved to the final fasta database (D), while entries where homologues were found in the subject files are indexed for further processing. (C) For each homologue group, a reference sequence is chosen, and polymorphic peptides are added into a customized entry (“Artificial Seq”), both being copied into the final database. (E) The Query file is removed from the dataset, the remaining subject files will have entries removed (proteins which were already defined as a homologue of a Query protein), and a new fasta file is selected as Query for a new BLASTP round. Such rounds will be performed until all files are analyzed.

Regarding all indexed homologues, our script then defines the longest version of the protein as reference (not necessarily from same strain used as query for the BHH step above). The reference sequence is copied integrally into the partial version of the final output. The script performs an *in silico* trypsin digest of the protein sequences, and compare amino acid composition of all generated tryptic peptides. Peptides shorter than 7 or longer than 35 amino acids are excluded. Whenever different tryptic peptide sequences are observed in the non-reference dataset (due to differences of TSS choices, SAAs or indels which result in frame shifts), they are added into the final output under an artificially created protein entry (Omasits et al., 2017). Amino acid changes result from poorly annotated sequences, where X or U are present in the sequence as an amino acid, are not considered. Finally, a log file is created, reporting and classifying all differences observed, describing the amino acid changes and also in which strains those were observed.

### Database analysis – Experimental design and statistical rationale

To investigate how each dataset contributes to the final database size, for each species we constructed databases using 5, 10, 15, 30 and 65 randomly selected strains. This was performed a total of three times to decrease the chances that a randomly selected strain with very unique features might interfere with the final result. Each database had its MS search space size measured regarding number of entries and number of unique tryptic peptide sequences. Pangenomic size calculations were based on the panX tool (Ding et al., 2018).

### Mycobacterial Cell Culturing

Stock cultures of Mtb strains TB179, TB861, TB1430, TB1593, TB1659, TB1841, TB1945 and TB2666 were inoculated into mycobacterial growth indicator tubes and incubated until positive growth was detected using the Bactec 460 TB system (BD Biosciences). Approximately 0.2 mL was inoculated onto Löwenstein-Jensen medium and incubated over 6 weeks with weekly aeration until colony formation. Colonies were transferred into 20 mL of supplemented 7H9 Middlebrook medium (BD Biosciences) containing 0.2% (v/v) glycerol (Merck Laboratories), 0.1% Tween 80 (Merck Laboratories), and 10% dextrose, catalase. Once the culture reached an A600 of 0.9, one mL was inoculated into 80 mL of supplemented 7H9 Middlebrook medium and incubated until an A600 between 0.6 and 0.7 was reached. All steps were performed at 37 °C until Mtb cells in mid-log growth phase were used for whole cell lysate protein extraction.

### Preparation of Crude Mycobacterial Extracts

Mycobacterial cells were collected by centrifugation (10 min at 2500 × g) at 4°C and resuspended in 1 mL of cold lysis buffer containing 10 mMTris-HCl, pH 7.4 (Merck Laboratories), 0.1% Tween 80 (Sigma-Aldrich), one tablet per 25 mL Complete protease inhibitor mixture (Roche Applied Science), and one tablet per 10 mL phosphatase inhibitor mixture (Roche Applied Science). Cells were transferred into 2 mL cryogenic tubes with O-rings, and the pellet was collected after centrifugation (5 min at 6000 × g) at 21 °C. An equal volume of 0.1 mm glass beads (Biospec Products Inc., Bartlesville, OK) was added to the pelleted cells. In addition, 300 μL of cold lysis buffer including 10 μL of 2 units/ml RNase-free DNase I (New England Biolabs) was added, and the cell walls were lysed mechanically by bead beating for 20 s in a Ribolyser (Bio101 Savant, Vista, CA) at a speed of 6.4. Thereafter the cells were cooled on ice for 1 min. The lysis procedure was repeated three times. The lysate was clarified by centrifugation (10,000 × g for 5 min) at 21 °C, and the supernatant containing the whole cell lysate proteins was retained. Thereafter the lysate was filter-sterilized through a 0.22 μm-pore Acrodisc 25 mm PF syringe sterile filter (PallLife Sciences, Pall Corp., Ann Arbor, MI), quantified using the Coomassie Plus Assay Kit (Pierce), and stored at − 80 °C until further analysis.

### Gel Electrophoresis and In-gel Trypsin Digestion

Each whole cell lysate (60 μg) were mixed with electrophoretic sample buffer (NuPAGE kit, Invitrogen, CA) containing 100 mM DTT, and heated for 5 min at 95 °C prior to the electrophoretic run. Proteins were separated in triplicate using a NuPage 4−12% Bis-Tris Gel in 2-N-morpholine ethane sulfonic acid (MES) buffer (Invitrogen) at 200 V for approximately 40 min. Proteins were visualized using a Colloidal Coomassie Novex kit (Invitrogen). After staining, each triplicate was cut into 15 fractions, sliced into smaller pieces and submitted to in gel digestion with trypsin, as previously described (Tomazella et al., 2012). Briefly, the proteins in the gel pieces were reduced using10 mM DTT for 1 h at 56 °C and alkylated with 55 mM iodoacetamide for 45 min at room temperature. The reduced and alkylated proteins were digested with 0.125 μg of trypsin (Sequence grade modified, Promega, WI, USA) for 16 h at 37 °C in 50 mM NH4HCO3, pH 8.0. The reaction was stopped by acidification with 2% trifluoracetic acid (Fluka, Buchs, Germany). The resulting peptide mixture was eluted from the gel slices, and further desalted using RP-C18 STAGE tips (Rappsilber et al., 2003). The peptide mixture was dissolved in 0.1% formic acid 5% acetonitrile for nano-HPLC-MS analysis.

### LC-MS/MS analysis

Peptides were separated by reversed-phase chromatography in an Acclaim PepMap 100 column (C18, 3 μm, 100 Å) (Dionex) capillary of 12 cm bed length and 100 μm inner diameter self-packedwith ReproSilPur C18-AQ (Dr. Maisch GmbH, Ammerbuch-Entringen, Germany), using a Dionex Ultimate 3000 nano-LC system connected to a linear quadrupole ion trap-Orbitrap (LTQ-Orbitrap) mass spectrometer (Thermo Electron, Bremen, Germany) equipped with a nanoelectrospray ion source. Peptides were onto loaded to the column with a flow rate of 0.3 mL/min of 7−40% solvent B in 87 min and then 40−80% solvent B in 8 min. Solvent A was aqueous 2% ACN in 0.1% formic acid, and solvent B was aqueous 90% ACN in 0.1% formic acid.

The mass spectrometer was operated in the data-dependent mode to automatically switch between Orbitrap-MS and LTQ-MS/MS acquisition. Survey full-scan MS spectra (from m/z 300 to 2000) were acquired in the Orbitrap with resolution of R= 60,000 at m/z 400 (after accumulation to a target of 1,000,000 charges in the LTQ). The method used allowed sequential isolation of the most intense ions (up to six, depending on signal intensity) for fragmentation on the linear ion trap using collisionally induced dissociation at a target value of 100,000 charges. For accurate mass measurements, the lock mass option was enabled in MS mode, and the polydimethylcyclosiloxane ions generated in the electrospray process from ambient air were used for internal recalibration during the analysis(Olsen et al., 2005). Target ions already selected for MS/MS were dynamically excluded for 60 s. General mass spectrometry conditions were as follows: electrospray voltage, 1.5 kV; no sheath or auxiliary gas flow. Ion selection threshold was 500 counts for MS/MS, an activation Q-value of 0.25 and activation time of 30 ms were also applied for MS/MS.

### MS Data analysis

Raw MS files of all Mtb and *S. aureus* datasets were analyzed by MaxQuant version 1.5.2.8. Peptides in MS/MS spectra were identified by the Andromeda search engine (Cox et al., 2011b). For Mtb dataset, MS files were searched against a database containing all sequences from 65 strains without any processing (263,683 entries), or the database generated by our tool using the same 65 strains (15,979 entries). The *S. aureus* dataset was searched against a database containing all sequences from 194 strains without any processing (523,281 entries) or the MSMSpdbb generated database (15,073 entries). All bacterial sequences were obtained from Genbank as written above (August 2017). MaxQuant analysis included an initial search with a precursor mass tolerance of 20 ppm which were used for mass recalibration (Cox et al., 2011a). Trypsin without proline restriction was used as enzyme specificity, with two allowed miscleavages and no unspecific cleavage allowed. In the main Andromeda search precursor mass and fragment mass were searched with initial mass tolerance of 6 ppm and 0.5 Da, respectively. The search included variable modifications of oxidation of methionine and protein N-terminal acetylation. Carbamidomethyl cysteine was added as a fixed modification. Minimal peptide length was set to 7 amino acids. The false discovery rate (FDR) was set to 0.01 for peptide and protein identifications, and since the aim is to compare database performance, no further score or PEP thresholds were used for individual MS spectra. In the case of identified peptides that are shared between two proteins, these are combined and reported as one protein group. Protein and peptide datasets were filtered to eliminate the identifications from the reverse database and from common contaminants.

## Results

### Implementation and database format

MSMSpdbb was originally designed to use gene assembly data from Genbank from selected bacterial strain genomes to construct protein sequence databases, which considered all possible sequence variations (SAP, TSS choice, for example) on a non-redundant manner (i.e., without extensive entry usage for very similar sequences). At the current version we modified it to perform BBH BLAST, which is an approach more used to define orthologous sequences, rather than Cluster BLAST. Database formatting was also modified; non-reference peptides are now inserted as new entries in the database.

On average, the assemblies used on this work had a number of genes ranging from 910 in *Chlamydia trachomatis* (average value to all strains) to 6092 in *Pseudomonas aeruginosa.* The analyzed species were selected not solely due to a higher number of strains with complete genome sequences available but also considering the complexity of their pangenomes. Nine of the selected species have pangenomes that can be very diverse (for example, *Escherichia coli*, 2,459 core genes to approximately 26,000 pangenomic genes) or just a fraction of the core genome (such as *Chlamydia trachomatis* and Mtb, which have only one accessory gene to every five core genes) (Rasko et al., 2008;McInerney et al., 2017;Ding et al., 2018).

Figure 1 illustrates the approach workflow: briefly, a query strain is selected, and a pair wise comparison is performed with the remaining subject strains. Unique entries from the query strain (i.e., sequences with no significant hit to any other protein entries in subject strains) are then saved in the final database. Protein sequences across strains that share the required levels of identity and similarity are further processed. The longest version is then used as a reference (regardless of which strain it is originated from) and saved into the final database. Tryptic peptides between reference and non-reference sequences are compared. When a tryptic peptide from a non-reference sequence do not match any of the reference sequence (due to a different TSS choice or a SNP for example), a new protein entry is created in the final database. This new entry will contain all non-reference tryptic peptides that do not match the reference sequence. This process will be repeated until no strains are left to be used as query, or until no protein entries are left in yet-to-be selected query strains. A log file is also created, reporting all entries that were matched and classifying all tryptic peptides which were added to the final database, showing the type of event (TSS choice, SNP etc) and in case of SNP, the resulting amino acid substitution.

Figure 2 illustrates a typical homologue comparison leading to the detection of polymorphic peptides, which will be added to the database as an artificial entry. The hypothetical protein AEM98552 (Rv0104 on the Tuberculist annotation (Lew et al., 2011)) was set as reference, and 9 other different variants were observed. Those variants are the result of the different combinations including: two additional TSS choices at position 9 and 118; three SAPs at positions 402 (Tyr → His), 477 (Gly→ Arg) and 493 (Leu→ Val); a not-classified genetic modification that leads to alternate amino acid sequence after position 475 (yellow box, named as frameshift/indel); and finally, two polymorphisms leading to premature stop codons and generating different c-terminal tryptic peptides ending at position 378 and 426. For example, the variant from strain CAS/NITR204 (AGL25567) contains two SAPs, while the variant from strain CCDC5180 (AEJ48929) has a different TSS choice at position 9 of reference, and a premature stop codon, truncating the protein at position 378. Unique tryptic peptides characterizing all such polymorphisms are added in the database as an additional entry.

**Figure 2.**
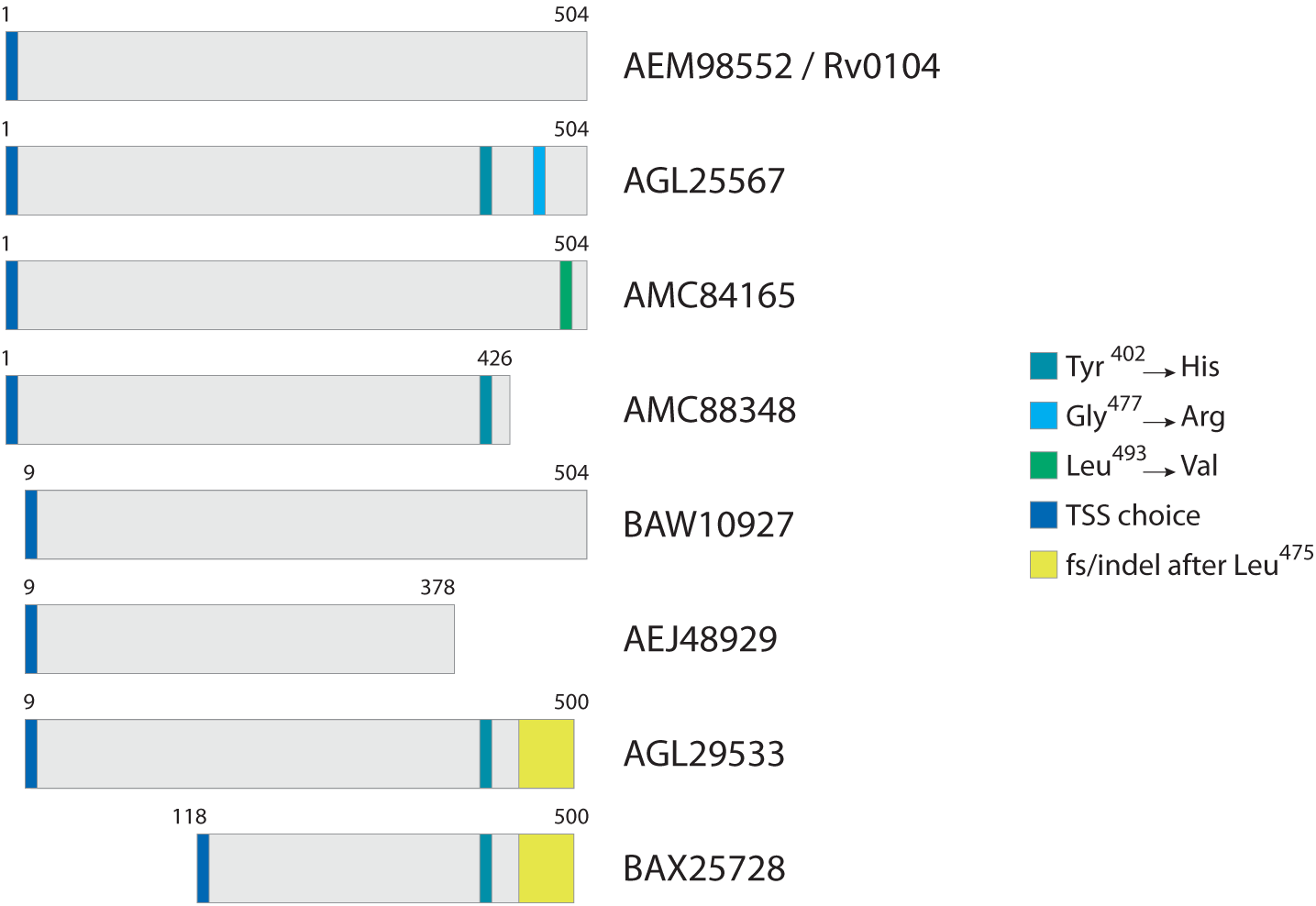
Typical homologue comparison. For protein Rv0104, in this case selected as reference, three SAPs were identified across all Mtb strains, in addition to two TSS choices, two premature stop codons, and a complete amino acid sequence change (region in yellow) most probably from an indel.

### Database size increase ratio per species

Databases were created for ten species to measure if the rate of increase in the database size could be impacted by unique features of each genome, such as the size of a pangenome. For a fair comparison among species, we built databases using a maximum of 65 strains, since at the time Mtb had only 65 strains with complete sequenced genomes available. To assist the visualization of how the database size is incremented as more strains are used, we also created databases using 5, 10, 20 and 30 randomly selected strains of each species. And to avoid outlier behavior in casea strain with very unique annotation being randomly selected, every database was created three times in total. The rand.pl script provided with the tool performs all such steps automatically, if the data needs to be reproduced. We then counted total number of protein entries and tryptic peptides in each database.

Supplementary Table 1 shows all values measured for all tests including: number of proteins (Supplementary Figure S1), number of annotated proteins (excluding artificial entries created to accommodate polymorphisms) (Figure 3 and Table I), and number of peptides with or without miscleavages allowed (Supplementary Figure S2). Figure 3 shows graphically the increase in number of annotated proteins in all species, and Table I describe number of annotated proteins in the database using 65 strains, rate of database size increase, number of annotated proteins added per strain on average, and size of pangenome for each species. As expected, the increase rate in the number of annotated proteins in the final database correlated with the size of the gene pool in the pangenome. *E. coli*, which has the largest genetic pool described, had indeed the larger database increase rate of all species, at 4.6x increase compared to the average number of genes per strain. The three exceptions for this were: *B. pseudomallei/mallei* with the 2^nd^ increase rate (3.53x) even though the pangenome size of *B. pseudomallei* is the 4^th^ most complex (no data is available regarding *B. mallei* pangenome); Mtb, which has the 2^nd^ less complex pangenome size, representing only 21% of total genetic pool, yet it has 4^th^ database increase rate (2.83x), surpassing *P. aeruginosa* which has a larger genome size and a bigger genetic pool; and *B. pertussis*, with no pangenome described so far and yet with a database increase rate (2.46x) which surpassed three species with characterized pangenomes.

**Figure 3.**
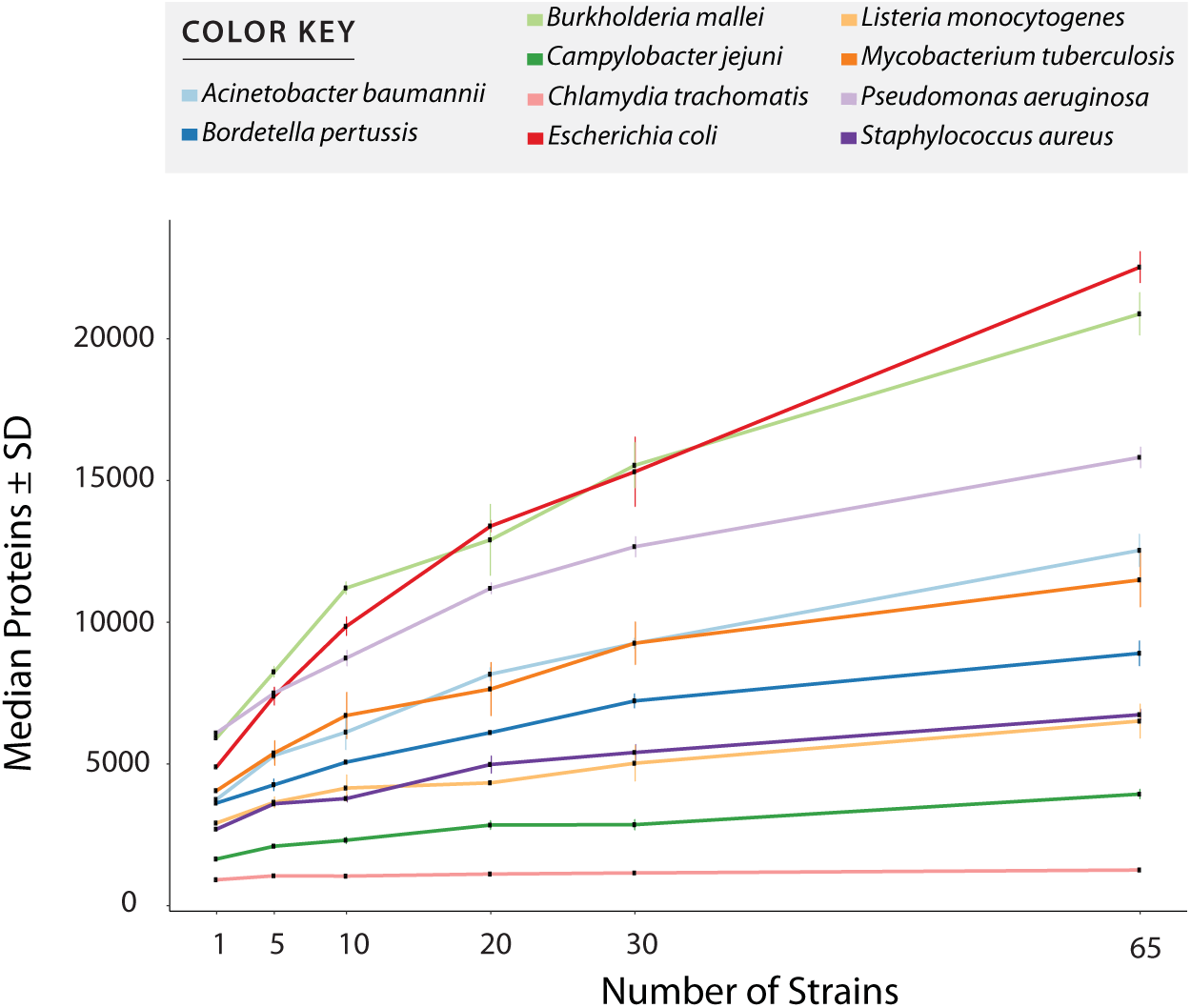
Number of entries per database after redundancy removal. The graph shows data when only considering reference and uniquely annotated sequences in all strains, i.e., customized entries containing polymorphic peptides where not counted. Values in x axis show the number of strains used for database creation. All values plotted in the graph are given in Supplementary Table 1.

**Table I.**
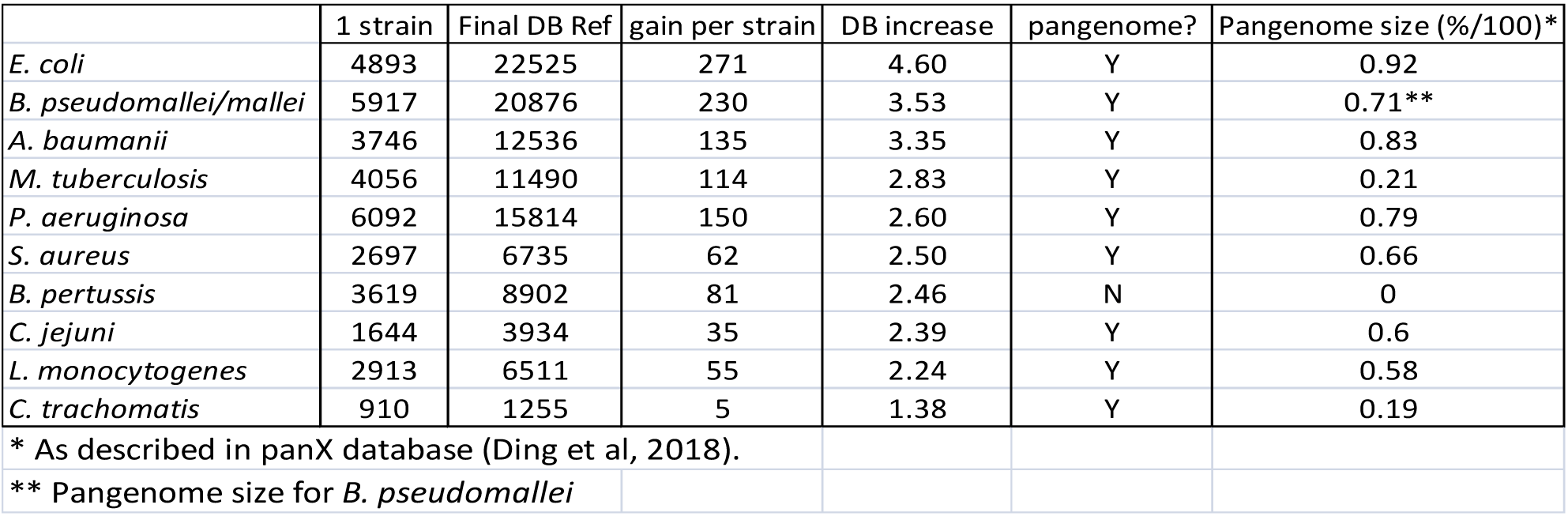
Database size increase and pangenome size in ten selected species

|  | 1 strain | Final DB Ref | gain per strain | DB increase | pangenome? | Pangenome size (%/100)* |
| --- | --- | --- | --- | --- | --- | --- |
| *E. coli* | 4893 | 22525 | 271 | 4.60 | Y | 0.92 |
| *B. pseudomallei/mallei* | 5917 | 20876 | 230 | 3.53 | Y | 0.71** |
| *A. baumannii* | 3746 | 12536 | 135 | 3.35 | Y | 0.83 |
| *M. tuberculosis* | 4056 | 11490 | 114 | 2.83 | Y | 0.21 |
| *P. aeruginosa* | 6092 | 15814 | 150 | 2.60 | Y | 0.79 |
| *S. aureus* | 2697 | 6735 | 62 | 2.50 | Y | 0.66 |
| *B. pertussis* | 3619 | 8902 | 81 | 2.46 | N | 0 |
| *C. jejuni* | 1644 | 3934 | 35 | 2.39 | Y | 0.6 |
| *L. monocytogenes* | 2913 | 6511 | 55 | 2.24 | Y | 0.58 |
| *C. trachomatis* | 910 | 1255 | 5 | 1.38 | Y | 0.19 |
| * As described in panX database (Ding et al, 2018). |  |  |  |  |  |  |
| ** Pangenome size for *B. pseudomallei* |  |  |  |  |  |  |

### Proteomic performance in diverse MS datasets

When we first developed the MSMSpdbb approach (de Souza et al., 2010), the number of genomic information available was a fraction of the amount currently in existence. For example, the analysis validation performed on an Mtb dataset used a database with only 8 strains (5 from *M. tuberculosis* and 3 from *M. bovis).* We then wondered if the approach would still be valuable as hundreds of genome information is available to a single species. This was tested by performing proteomic analysis in two independent datasets.

The MSMSpdbb processed database created by the tool contained 15,996 (Mtb), 15,073 (*S. aureus*) protein entries. In comparison, merely concatenated databases using all sequences from the available strains for each species created a database with 263,683 protein entries for Mtb and 523,281 protein entries for *S. aureus.* From now on, all MSMSpdbb processed databases are called DB1, and simply concatenated databases are named DB2 for simplicity. The MS datasets were then challenged for peptide identification using the respective DB1 and DB2 databases for each species. It is important to note that all theoretical unique tryptic peptides present in DB2 are also present in DB1 (data not show). This means that MSMSpdbb did not lose peptide sequences during its database construction, and that the differences observed below regarding number of identified peptides is not due to missing information in DB1. The complete results folders from each MaxQuant search including protein and peptide lists with relevant identification parameters are provided (see Data availability section).

In all datasets, when comparing the results from DB1 and DB2 searches, the number of peptides identified was very similar in both searches (Figure 4). For example, the Mtb dataset identified 32,986 unique peptides in both searches, while 515 peptides were identified only in the DB1 search and 1,116 peptides only in the DB2 search result (Figure 4A). When all peptides are considered, 95.28% of the peptides were identified in both searches. For *S. aureus*, DB1 and DB2 searches were in agreement for 92% of the identified peptides in all searches combined. We observed that the majority of such database exclusive identifications are distributed in the lower range of the spectra scoring. For all datasets, the median MaxQuant score for peptides identified in only DB1 and DB2 were in the range of 52 to 75, while the median scores of peptides identified both searches was from 128-137 (see histograms Figure 4). If a MaxQuant posterior error probability (PEP) score lower than 0.01 is also considered together with regular MS2 score, the numbers of exclusive peptides drop by half in DB1 and DB2 for all datasets considered (data not show).

**Figure 4.**
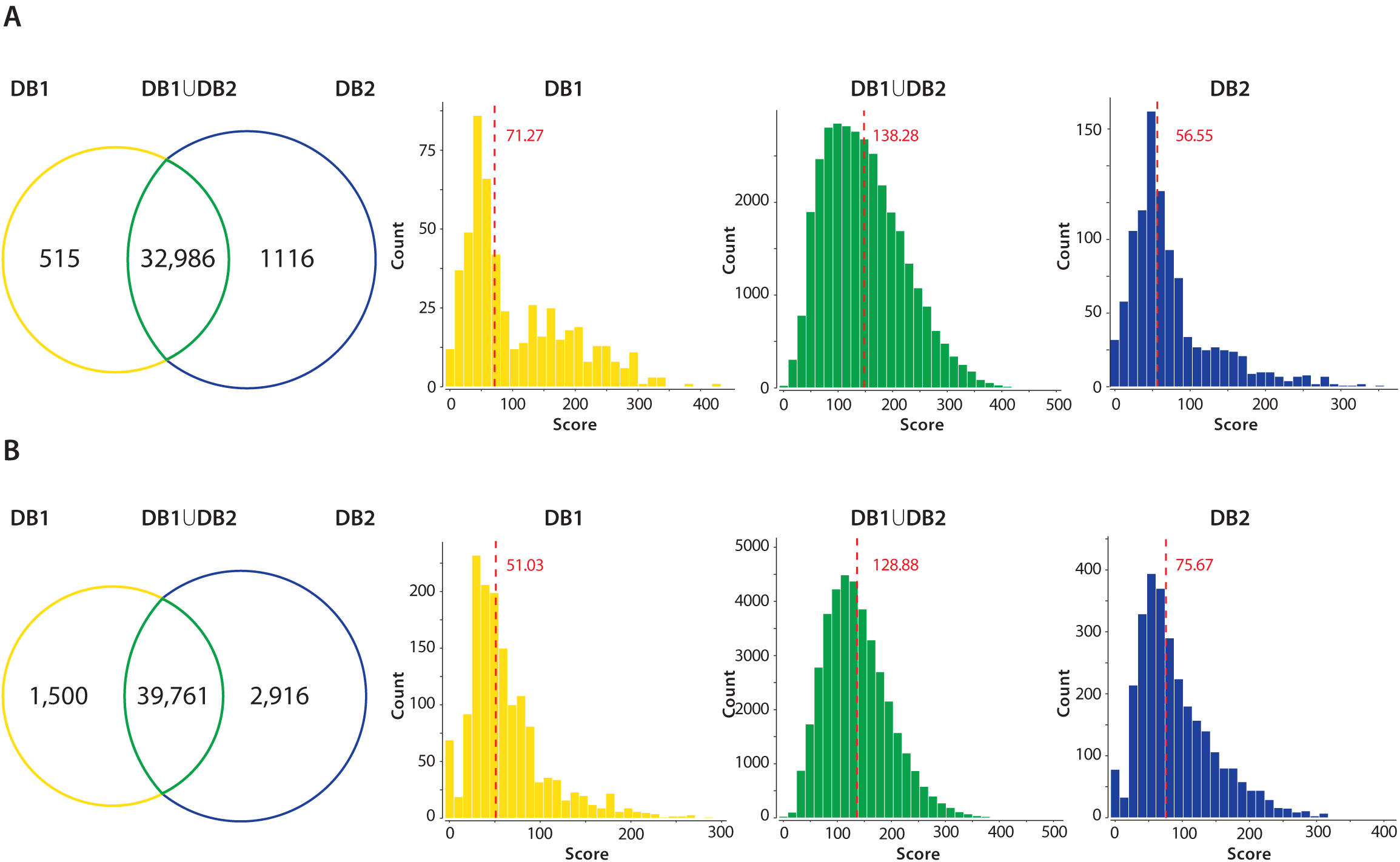
DB1 vs. DB2 peptide identification performance. All unique tryptic peptides identified in DB1 and DB2 in the Mtb (A) and *S. aureus* (B) datasets are compared. Venn diagrams show that majority of tryptic peptides were identified in both searches. Peptide scores of each group were plotted in a histogram, and a median was calculated. Median of the score distribution for peptides exclusive to DB1 or DB2 ranged from 51.03 to 75.67, while median scores of peptides identified on both searches were 138.28 for Mtb and 128.88 for *S. aureus*.

Multiple TSS polymorphisms for the same protein were also observed even for the same strains. Figure S3 shows the position of three peptides; all predicted as possible N-terminal peptides of the protein lipase lipV. MIIDLHVQR (Score 93.4, PEP 0.0019, Figure S3B) represents the most 5’ prediction; LTIHGVTEHGR (Score 85.9, PEP 0.0005, Figure S3C) starts at position 19 of reference protein and HGVTEHGR (Score 100, PEP 0.0077, Figure S3D) starts at position 22 of reference. The longer form is the most abundant in almost all samples, except S1430 where only LTIHGVTEHGR was identified and S2666 where the N-terminal HGVTEHGR is slightly overrepresented than MIIDLHVQR. Finally, the DB1 results also included 5 polymorphic peptides from the artificial entries, which could not be found in the DB2 database. Closer inspection of these peptides revealed that they all contain missing cleavage sites. For example, the identification of peptide GEGAAVVVIKEGSIGGGTVFVYSGR from gene PpsC includes two polymorphic peptides that were concatenated in DB1, GEGAAVVVIK corresponding to positions 274-283 in the reference protein, and EGSIGGGTVFVYSGR corresponding to positions 564-578. Clearly, the artificial entries in this case contain two concatenated polymorphic peptides which are not adjacent in the reference protein. The fact that we accept miscleavages as a parameter during the search allows a clearly false-positive identification for artificial entries. The spectra and identification score for this concatenated peptide is given in Supplementary Figure S7, and as expected, the identified spectra has poor score and sequence coverage. Supplementary Table 2 lists all five false-positive peptides.

### Database size impact in computational performance

As shown before, database size reduction from DB2 to DB1 was approximately 16x to Mtb and 35x to *S. aureus.* By inspecting MaxQuant log folder we calculated the amount of minutes used for each step in its proteomics pipeline. For Mtb, the time spent for the whole pipeline was reduced by half, from 853 minutes for DB2 to 434 minutes to DB1 (Figure 5). Main differences were observed for steps related to the peptide search, i.e., the ‘main search’ (5x) and the ‘second peptide search’ (4x) steps, and during the creation of results and other final files named ‘writing tables’ (3.5x). For *S. aureus* the differences were far more evident, as the complete pipeline took 7,553 minutes for DB2 and 2,147 minutes for DB1. Time reduction in peptide search steps were similar to Mtb at 4.5x and 4x respectively. But in addition to those, other steps which are related to read and finish the main and the second search results had improvements of 42 and 19.6 times respectively. Steps related to assignment and assembly of protein groups had improvements of 42 times, and ‘writing tables’ had a time improvement of 10.4 times.

**Figure 5.**
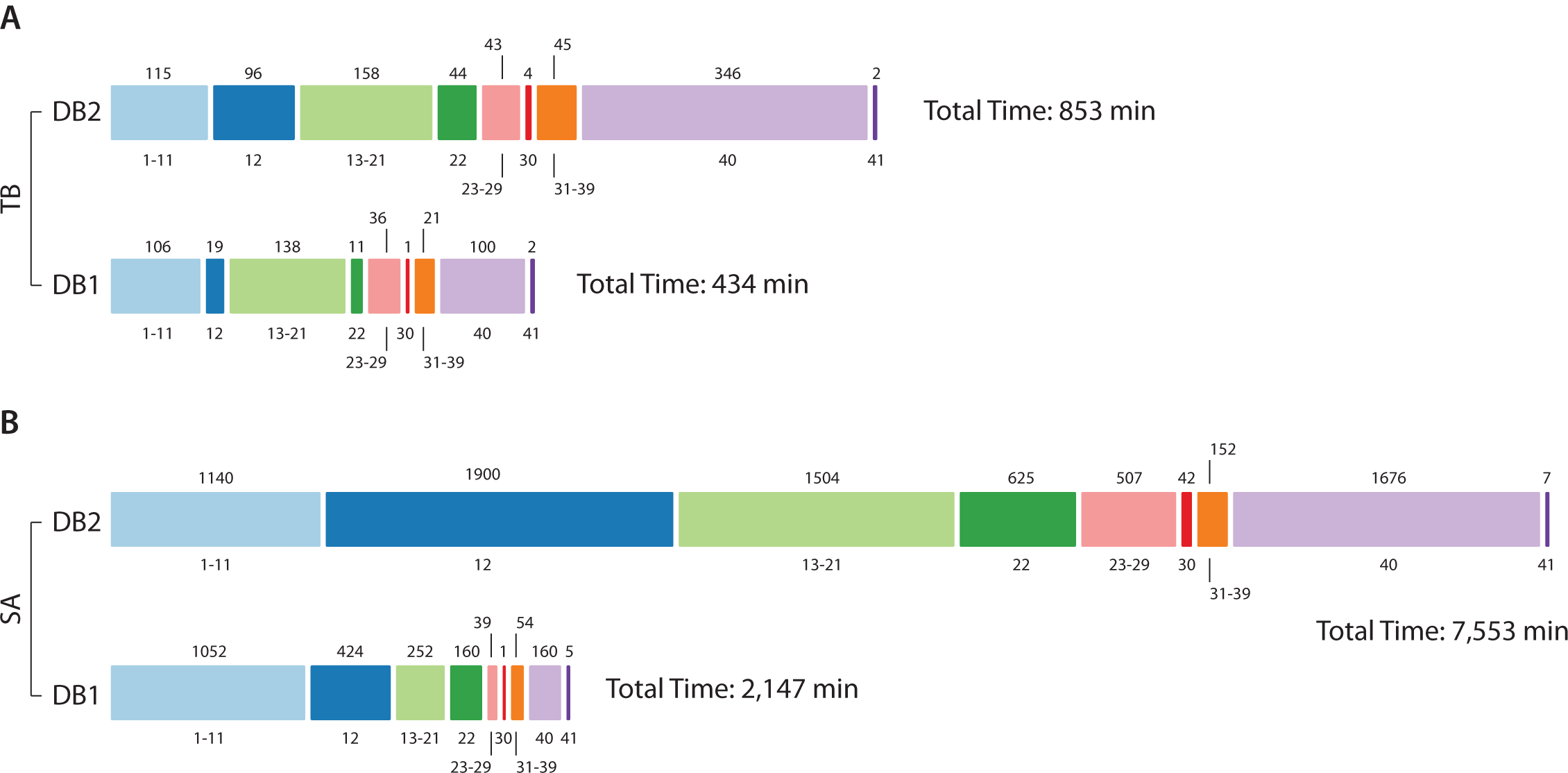
DB1 vs DB2 computational demand. Processing times for Mtb (A) and *S. aureus* (B) datasets. The bar is divided into MaxQuant stage number, as described in (Neuhauser et al., 2013) (value at bottom) and time in minutes is at the top of the bar. For Mtb, analysis was hasten mainly at stage 12 (Main search), 22 (Second peptide search) and 40 (Writing tables); in addition to those, for *S. aureus* improvements were also observed for post-search steps 13-21 and 23-29, and step 30 (“Protein groups assembly”).

## Discussion

Database size is a known bottleneck in proteomics, and its implications had been already discussed in proteogenomics and metaproteomics research (Jagtap et al., 2013;Heyer et al., 2017). Therefore, controlling database information is of key importance. A manner to achieve this is to construct customized databases where identical sequences/tryptic peptides from homologue proteins are not exhaustively present in different entries within the database. We had already applied this for the characterization of samples with unknown genetic background such as bacterial clinical strains (de Souza et al., 2011;Tomazella et al., 2012). Such databases allow the validation of coding regions and confirmation of sequence polymorphisms normally omitted from reference databases. This is a critical characterization, considering that even genomes with two decades of considerable investigation still have gene annotation issues (Abascal et al., 2018). It is true even for prokaryotes (de Souza et al., 2008;de Souza et al., 2009), with their simpler genomic structure and higher coding density. However, as the amount of genomic information exponentially increases with time, we wished to further evaluate the impact of such approach with the current amount of available genomic data.

First, we were curious to investigate how different species with diverse genetic features (such as the presence of a pangenome) would impact database size and structure. As expected, database size increase correlated with the pangenome complexity of each species. Surprisingly Mtb and *B. pertussis* databases size increased at rates higher than expected based on the low complexity of their core and pangenomic genes. Previous data from Mtb suggested that database size increment was not heavy, when at the time only five Mtb strains and three strains of *M. bovis* were considered (de Souza et al., 2011). Our data here however shows that Mtb database size increment was one of the most prominent, arguably the most relevant if its small pangenome is considered. This suggests that as additional strains were sequenced, variations in gene annotation performed were more acute then observed for the other nine species. Something similar could be assumed about *B. pertusis* databases, which has no pangenome and database increase was higher than three species with characterized pangenomes. However, *C. trachomatis, L. monocytogenes* and *C. jejuni* have smaller genomes than compared to *B. pertussis*, all also having higher density of coding regions (Parkhill et al., 2000;Parkhill et al., 2003;Thomson et al., 2008). More nucleotide regions in the genome marked as non-coding will offer more opportunity for different strategies to generate conflicting gene annotation data, which could explain *B. pertussis* database size increase in our analysis. The database size increase for *Burkholderia pseudomallei* and *Burkholderia mallei* was not taken into consideration by us on this analysis, because while we decided to merge both species as one based on genotyping data (Godoy et al., 2003), large scale rearrangements in *B. mallei* (Losada et al., 2010) might be interfering with our approach.

We then selected two MS datasets from Mtb and from *S. aureus* to identify peptides using a routine probabilistic-based approach, but now using the complete set of available genomes (194 strains) for *S. aureus.* Overall, DB1 and DB2 performance was very similar, and differences were negligible and mostly could be ignored to their low scoring nature. A minority of valid identifications exclusive to DB2 were N-terminal peptides containing miscleavages that were not selected by MSMSpdbb (i.e., the peptides without miscleavages were shorter than 7 amino acids and not present in the selected reference protein) and therefore were absent in DB1 due to the parameters used for DB1 construction. It is worth mentioning that surprisingly, we have identified multiple N-terminal predictions for the same protein, in some cases observed in the same strain (Supplementary Figure 3). While normally the identification of a protein N-terminal peptide in proteogenomics is used to validate and confirm the TSS of the protein, our data suggest that excluding the remaining TSS choices from the database might be undesirable, considering they might be later identified when additional strains are analyzed by MS.

The advantage of using a concatenated database is evident when computational performance is measured. The main steps in Andromeda/MaxQuant pipeline which involve Fasta reading (‘main’ and ‘second peptide’ searches) were similarly improved in both datasets tested. In *S. aureus* dataset, where DB1 concatenation is larger when compared to DB2 (from approximately 500,000 entries to 15,000 entries), time processing improvements were observed in additional stages of the pipeline. Particularly steps related to handle of peptide search engine results and assembly of the protein groups to be reported as final protein identifications. The step which creates the final output files (‘Writing tables’), even though do not directly use the Fasta file, was also significantly improved. This makes sense, as MaxQuant will report all entries from the database that share identical peptides. Entries from a protein with high sequence identity present in all 194 strains for *S. aureus*, for example, will have all 194 entries recorded in the output if that protein is identified. For DB1, since those 194 highly identical sequences are concatenated into a single reference entry, the output reporting is vastly simplified.

As mass spectrometry evolves and as detection sensitivity and peptide identification coverage improves, one of the last bottlenecks in metaproteomics research is database design. Building databases for metaproteomics is often achieved by two approaches: either by collecting metagenomic or metatranscriptomic data from the sample; or by concatenating protein sequences from a public repository such as Uniprot (for recent reviews see (Heyer et al., 2017;Starr et al., 2018)). Regarding the metagenomic approach, even with the advances of next-generation sequencing methods, reads and contigs assembly in complex samples is still challenging (Heyer et al., 2017). On the other hand, concatenating protein sequences from multiple organisms from a public repository results in very large databases, making the analysis time costly and susceptive to high false-positive rates (Tanca et al., 2013). False-positive issues had been mostly solved by database size restrictions using a multi-step peptide search or multiple peptide searches using smaller datasets of refined sequences on each search and further analysis to combine results (Jagtap et al., 2013;Muth et al., 2016;Zhang et al., 2016;Liao et al., 2018;Muth et al., 2018). While those are successful methods, they are mostly provided as complete pipelines. We believe that providing solutions solely at Fasta file level offers more freedom to researchers, as they can use whichever peptide search engines or pipelines they prefer. We predict that controlling database size such as done by MSMSpdbb without losing peptide identification coverage and drastically reducing computational processing time will be of key importance for metaproteomics.

### Data availability

Scripts are fully available at https://github.com/karlactm/Proteogenomics. All Mtb mass spectrometry files, and MaxQuant search parameters and results are available at ProteomeXchange with the accession number PXD011080. *S. aureus* mass spectrometry files are found in ProteomeXchange as PXD006483 (Hoegl et al., 2018) and PXD000702 (Depke et al., 2015).

## Acknowledgements

This work was supported by: grant 23038.004629/2014-19 to SJS and GAS from the Coordenação de Aperfeiçoamento de Pessoal do Ensino Superior; grants 305233/2015-7 to SJS and 406630/2016-0 to GAS from Conselho Nacional de DesenvolvimentoCientífico e Tecnológico (CNPq); and grants 175141 (HWG) and 183418 (HGW/RW) from the Norwegian Research Council. The manuscript has been released as a Pre-Print at bioRxiv with doi: https://doi.org/10T101/378117.

## Authors’ contribution

KCTM designed the tool and performed bioinformatic analysis; SF and GGT generated Mtb samples and proteomics data; RW, HGW, SJS and GAS supervised work and data analysis. GAS provided *S.aureus* proteomic analysis. KCTM and GAS designed experiments and analysis.

**Figure SI - Total number of entries per MSMSpdbbdatabase.** Values in x axis show the number of strains used for database creation. All values plotted in the graph are given in Supplementary Table 1.

**Figure S2 - Number of tryptic peptides per MSMSpdbbdatabase.** In (A) only fully tryptic peptides are considered, while in (B) peptides with up to two miscleavages are also counted, a standard search parameter choice in proteomics. Values in x axis show the number of strains used in the database. All values plotted in the graph are given in Supplementary Table 1.

**Figure S3 - Multiple TSS identified in protein lipV.** (A) Three possible TSS choices were identified in clinical Mtb strains. The most upstream prediction (MIIDLHNQR) is the most abundant variant observed in all strains, except strain S1430 which has only the variant with TSS at position 19 (ITIHGVTEHGR) of the reference sequence, and strain S2666 which TSS at position 22 (HGVTEHGR) is predominant. Two possible TSS choice variants were also observed at high levels for strain S1593. (B-D) MS2 spectra for all three peptides mentioned above.

**Figure S4 - False-positive identification in DB1.** MS2spectrum of identified peptide GEGAAVVVIKEGSIGGGTVFVYSGR, a concatenated sequence of peptide GEGAAVVVIK (pos 274-283 on PpsC protein) and peptide EGSIGGGTVFVYSGR (pos 564-578 on same protein). Note the low scoring of the peptide identification. All five false-positives identified due to concatenation of non-neighboring peptides are given in Supplementary Table 2.

