## Supplementary figures and images for "On the impact of the pangenome and annotation discrepancies while building protein sequence databases for bacteria proteogenomics"

### Figure S1

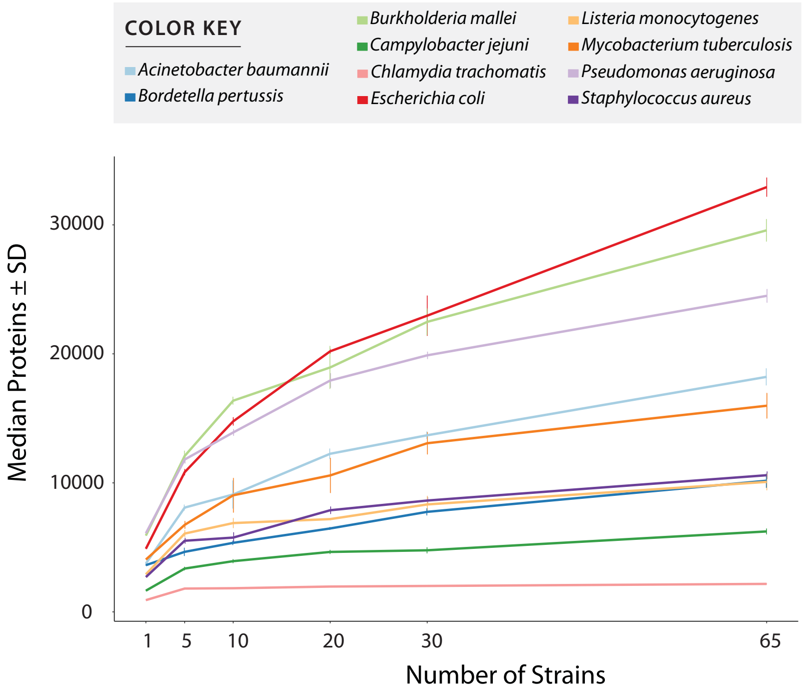

### Figure S2

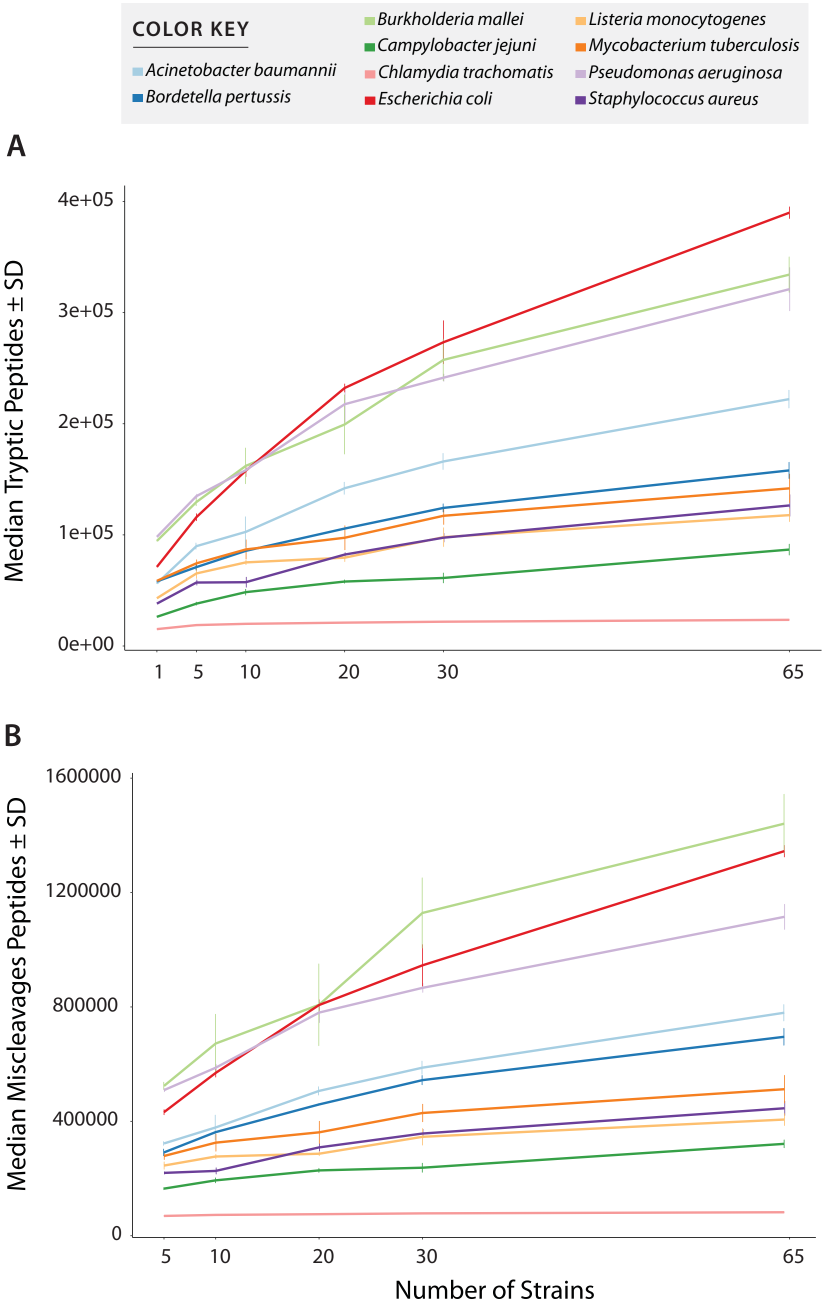

### Figure S3

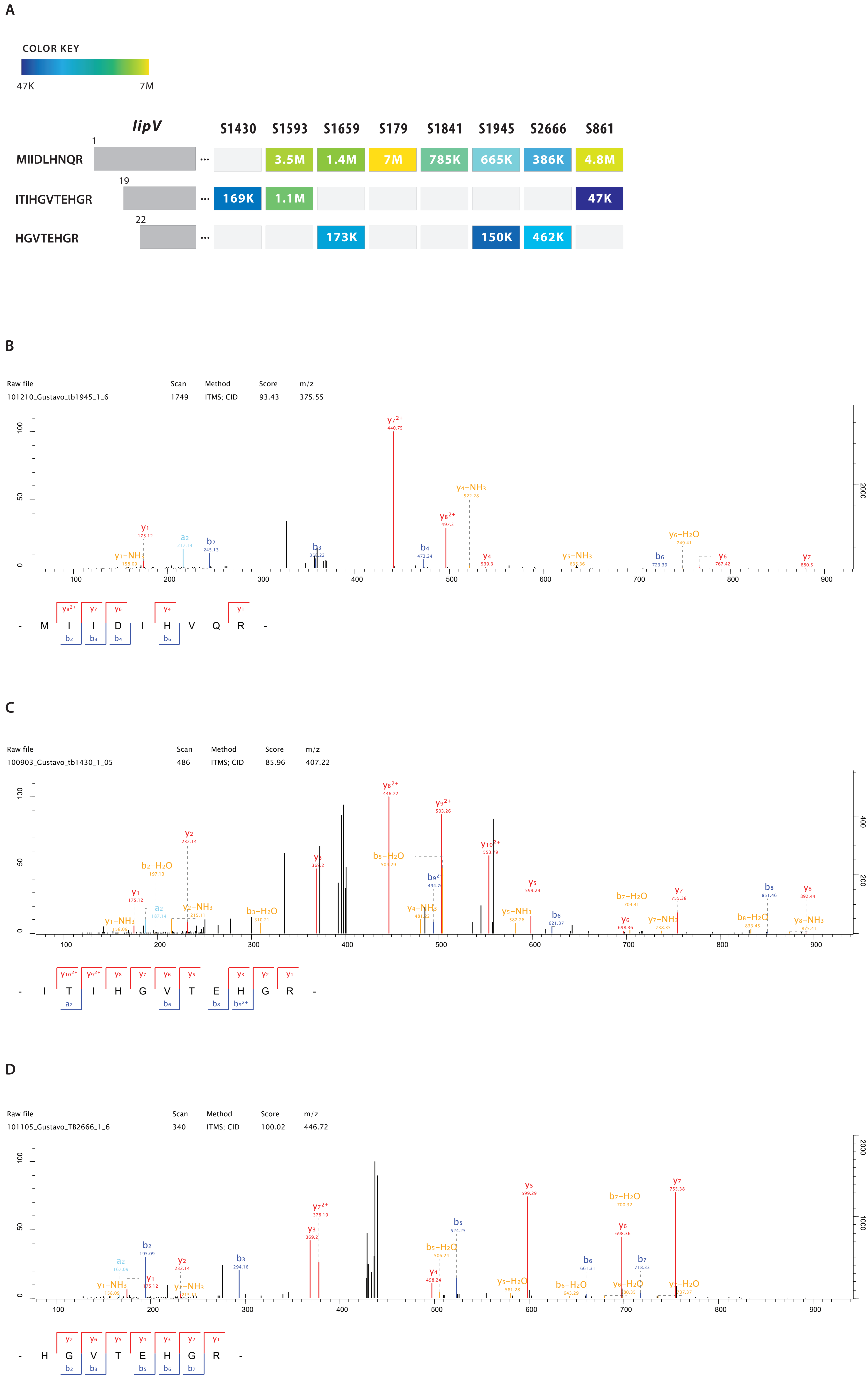

### Figure S4

| Raw file | Scan | Method | Score | m/z |
|---|---|---|---|---|
| 101230_Gustavo_1593_1_08 | 6739 | ITMS; CID | 27.55 | 804.11 |

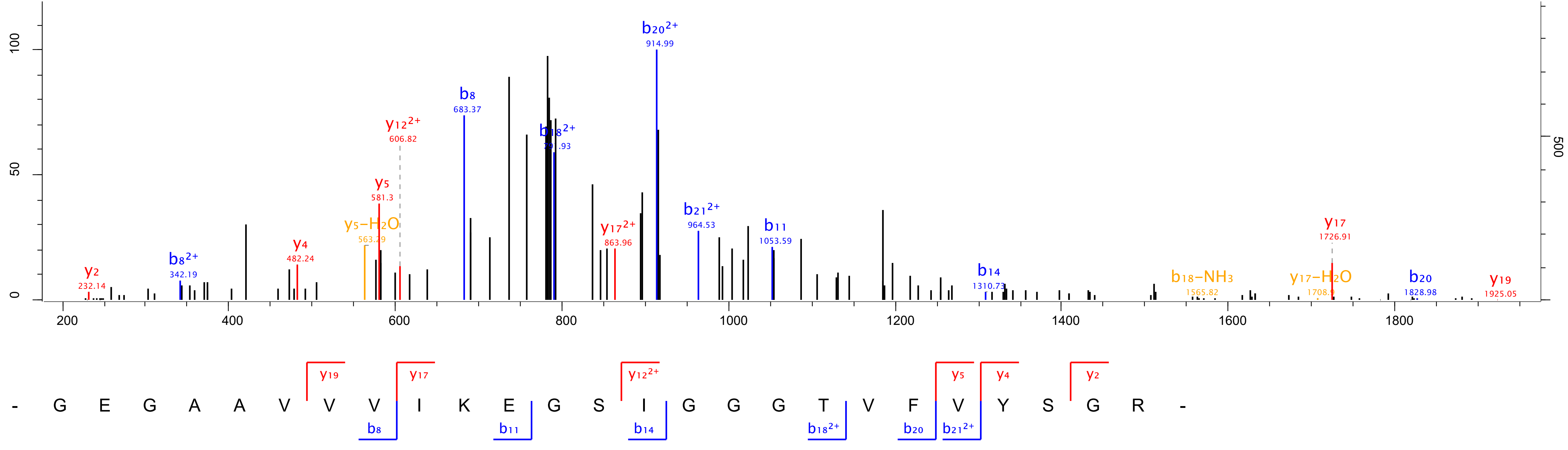
